# Deep learning for satellite image forecasting of vegetation greenness

**DOI:** 10.1101/2022.08.16.504173

**Authors:** Klaus-Rudolf Kladny, Marco Milanta, Oto Mraz, Koen Hufkens, Benjamin D. Stocker

## Abstract

The advent of abundant Earth observation data enables the development of novel predictive methods for forecasting climate impacts on the state and health of terrestrial ecosystems. Here, we target the spatial and temporal variations of land surface reflectance and vegetation greenness, measuring the density of green vegetation and active foliage area, conditioned on current and past climate and the local topography. We train two alternative recurrent deep learning models that rely on convolutional layers for forecasting the spatially resolved deviation of surface reflectance across a heterogeneous landscape from a specified initial state (*Baseline Framework*). We demonstrate efficiency of the Baseline Framework with respect to training convergence speed. Using data from diverse ecosystems and land cover types across Europe and following a standardized model evaluation framework (*EarthNet2021 Challenge*), results indicate increased performance in predicting surface greenness during drought events of the models presented here, compared to currently published benchmarks. Our results demonstrate how deep learning methods enable early-warning of vegetation responses to the impacts of climatic extreme events, such as the drought-related loss of green foliage.

## 1 Introduction

Recent hot and dry summers in Central Europe led to measurable and visible impacts on the functioning and structure of forests [1, 2, 3]. Combined heat and drought stress caused wide-spread premature canopy defoliation, tree mortality, and carbon (C) losses from ecosystems [2, 3, 4]. The timings and locations of such extreme event impacts can be identified from space as anomalously low vegetation greenness (*browning*), measured by satellite remote sensing of the surface reflectance at high resolution and with global coverage [5, 6]. Although such Earth observation data has yielded a wealth of information covering past climatic extreme events and vegetation responses, thresholds at which lasting impacts are triggered are difficult to anticipate and are often identified in retrospect [1, 2, 6]. Observed relationships of impacts and climate, including its history, can inform predictions that are based on modelling their functional relationships and thus forecast where and when impacts of unfolding meteorological extremes occur.

Large volumes of Earth observation and climate re-analysis data, combined with detailed information about the local conditions (soil, topography, land cover type) provide an opportunity to develop data-driven predictive models of impacts caused by extreme heat and drought. However, suitable machine learning algorithms have to be tailored to learn key factors that drive impacts by climatic extreme events across the landscape and over the seasons and model the distinct temporal dependencies and spatial heterogeneity of relationships.

Temporal dependencies arise because impacts of climate extremes and browning depend on the climate of preceding months. A progressive depletion of plant-available water stores evolves over several weeks [7]. Hence, for example, a dry spring can amplify the sensitivity of vegetation to hot and dry weather in summer. Furthermore, an early start of the season (early leaf unfolding dates) can enhance water losses during spring and thus lead to an early onset of water-stressed conditions in summer [8]. Similarly, favourable growth conditions in the early season can enable trees to develop a large total foliage area, making them sensitive to excessive water loss during dry conditions in the later season [9]. Hence, effective models must learn the temporal dependencies of multiple co-varying drivers.

Spatial heterogeneity arises because environmental conditions vary substantially across elevation, position along the hillslope (ridge vs. valley bottom), exposition (north vs. south), or upstream drainage area [10], and with small-scale variations in soil properties. Highly localized growth conditions and microclimates interact with variations in ecosystem properties to determine drought and heat impacts. Varying soil and plant rooting depth [11], vegetation access to water stored in weathered bedrock [12] and groundwater [13] and large variations in incoming radiation across different positions in the landscape drive large spatial variations in water stress [11]. This large spatial heterogeneity of impacts across the landscape (100 m - 1 km), combined with the fact that climate reanalysis data is commonly provided at much lower resolutions (10 - 100 km), poses a challenge for reliable predictions of impacts.

Potentially suitable machine learning model architectures have been developed for related tasks and may be applied for learning the distinct temporal dependencies and spatial heterogeneity of vegetation greenness anomalies in response to climatic extremes. In particular, a combination of convolutional and Long-Short Term Memory (LSTM) cells in deep neural network architectures have been shown to perform well on video prediction tasks [14, 15, 16] and may be repurposed for effectively forecasting the near-term evolution of satellite images, conditioned on the evolution of climate and the position in the landscape.

The high demand for reliable extreme events’ impact prediction and early warning, combined with the availability of large data volumes and the development of powerful machine learning algorithms, gave rise to EarthNet2021 - a formalized prediction challenge for satellite image forecasting [17]. EarthNet2021 provides Sentinel 2 satellite data for surface reflectance [18] and spatially aligned topography information, as well as temporally and spatially aligned climate re-analysis data [19]. The EarthNet2021 Challenge also defines a common training and testing framework and a unified model evaluation metric, enabling a standardized comparison and benchmarking of different models, i.e., competing submission to the challenge. Here, we implemented two alternative deep neural networks and show, using the EarthNet2021 data and their model evaluation framework, that both models are well-suited for the drought impact prediction challenge at hand.

## 2 Methods

### 2.1 Prediction task

We followed the prediction task defined by the EarthNet2021 Challenge [17]. As illustrated in Fig. 1, the task is to predict the future evolution of (high-resolution) surface reflectance, given its past evolution, given past and future (mesoscale) climate, and given high-resolution information of (time-invariant) topography. Datacubes (see also Sec. 2.2) of remotely sensed surface reflectance contain ten frames (or images, i.e., data arrays in longitude and latitude) from past and current time steps *t* = 1, … *T*_1_ as *context* and twenty frames for time steps *t* = *T*_1_ + 1, …, *T*_2_ as *target*. Climate data for all time steps and time-invariant topography information are used as model inputs for past and present time steps and guide predictions for future time steps. Datacubes are divided between a set used for model training and four distinct sets for testing, as defined by EarthNet2021 (see Sec. 2.4).

**Figure 1.**
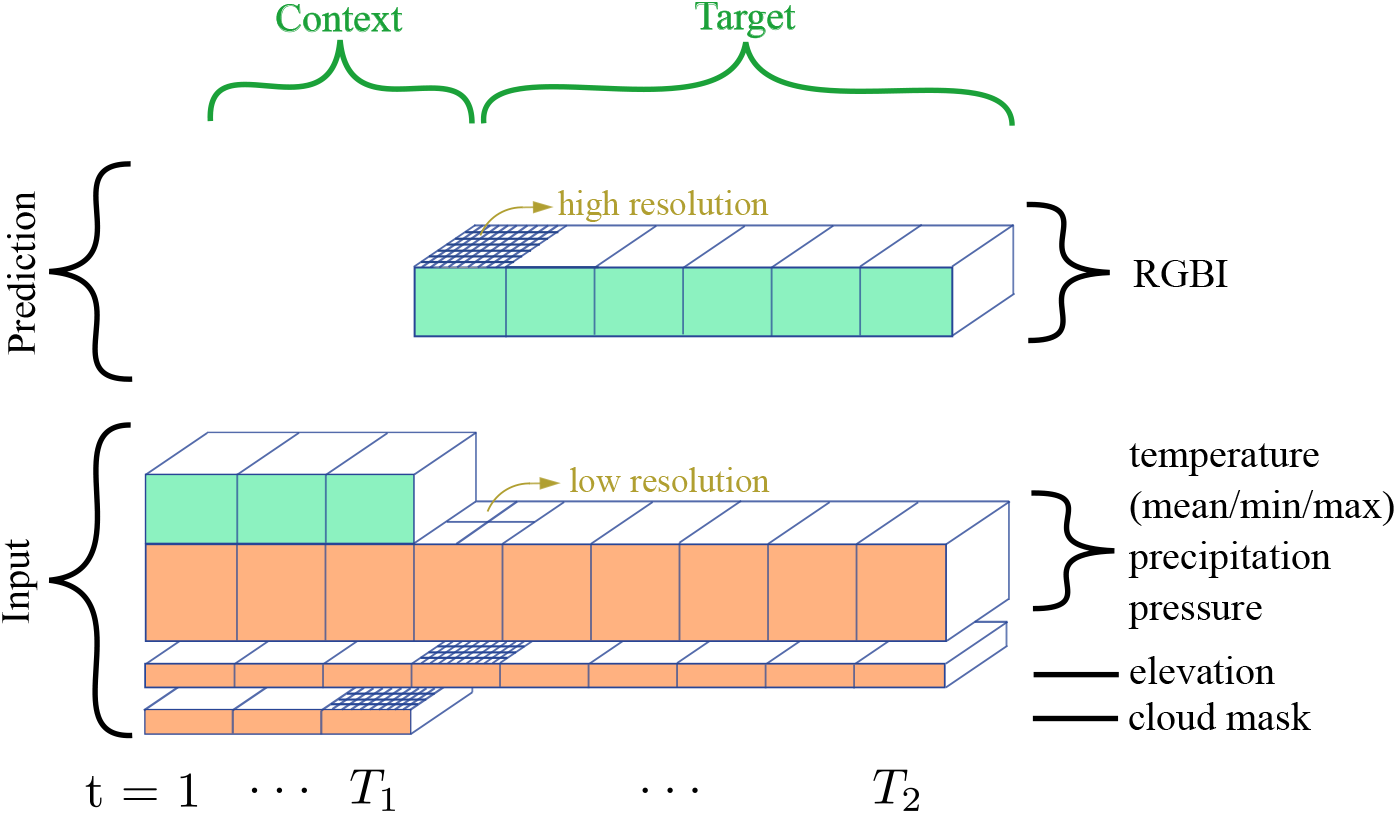
Illustration of the prediction task. Data is composed of *context* frames (past and current time steps *t* = 1, … *T*_1_), and *target* frames (future time steps *t* = *T*_1_ + 1, …, *T*_2_). High resolution remote sensing data of the surface reflectance in four spectral bands (’RGBI’ for red, green, blue, and infra-red) are used as model input for the context frames and are to be predicted as target for future time steps. Multiple low-resolution climate variables are used as model input for both the context and target time steps and thus guide predictions. Elevation from a digital elevation model is provided as a time-invariant model input for the context and the target time steps. The cloud mask for the context time steps specifies information in the RGBI frames that is to be ignored.

By targeting the prediction of future frames from past frames, the EarthNet2021 Challenge resembles a standard video prediction task. However, in contrast to the general video prediction framework, forecasting the motion of objects in the scene is not relevant here. In other words, the spatial arrangement of the land cover within a remote sensing data scene is largely constant and locomoting objects are absent or not relevant for the prediction task. The forecasting target is limited to the distinct evolution of land surface reflectance in separate, but fixed portions of the image. Furthermore, predictions are guided by the information of future climate, while climate-surface reflectance relationships are learned from their past dependencies and temporal dynamics. Additionally, time-invariant information about the topographic arrangement of the landscape is provided by the elevation map and may enable the learning of spatially varying climate-surface reflectance relationships, modified by the topographical setting. These features provide additional information for model predictions. We therefore refer to the task as a *strongly guided* (SG) video prediction task and adopt this term also for naming our models.

### 2.2 Data

The data used here were provided as part of the EarthNet2021 Challenge and consist of 23,904 training datacubes. Here, a datacube is a data array with two spatial dimensions, longitude and latitude, and a dimension in time *t*. Each datacube covers a geographical domain in longitude and latitude - a scene. Datacubes provided through EarthNet2021 are composed of high resolution data of remotely sensed surface reflectance, and of mesoscale resolution climate data.

Remote sensing datacubes represent scenes from the Sentinel 2 mission [18], covering a spatial extent of 128 × 128 pixels at 20 m resolution (2.56 × 2.56 km total extent), and providing surface reflectance every 5 days in four wavelength bands (corresponding to blue, green, red, and near-infrared light (RGBI)), complemented with a binary data quality mask defining the presence of clouds. Climate datacubes from E-OBS climate reanalysis data [20] are provided for each day. They have an extent of 80 × 80 pixels at a resolution of 1.28 km (corresponding to ≈ 0.1°, referred to as ‘mesoscale resolution’, covering 102.4 × 102.4 km total extent) and provide information on precipitation, sea level pressure, daily mean, minimum and maximum temperature. Additionally, time-invariant data layers of elevation from the EU-DEM digital elevation model [21] are provided at both high and mesoscale resolutions. Remote sensing and climate data cubes are spatially aligned such that within the geographical extent of a given climate data cube, multiple remote sensing data cubes are provided. A more detailed description of the data provided through EarthNet2021 is given by ref. [17].

In order to use the data for modelling here, we applied additional processing steps. The high-resolution elevation data were replicated for each time step. The mesoscale elevation data was not used. The daily meteorological data were aggregated to 5-day intervals, matching the frequency of the remote sensing data. From the original daily mean temperature, daily total precipitation and daily mean atmospheric pressure, we computed the mean across respective intervals of five days. For the daily minimum and maximum temperature, we took the minimum and maximum values across the 5-day interval, respectively. We used only the subset of climate data, matching the spatial extent of the remote sensing data cubes. Climate data outside this domain was not considered. Due to the presence of clouds, data completeness within the individual datacubes varied strongly. We discarded three datacubes from the training set that were affected by cloud-contamination of a subset of pixels throughout the entire context period.

### 2.3 Model

To address the temporal and spatial dependencies of the data and the prediction task, we used two variants of the Convolutional Long Short-Term Memory (ConvLSTM) network. The ConvLSTM is a convolutional adaptation of the standard LSTM [22] and is designed for the purpose of processing sequential image data - suitable for the spatio-temporal prediction task at hand. LSTMs are a subclass of recurrent neural networks, chosen here to satisfy our prior assumption about the task that the time dimension is shift-invariant. Recurrent neural networks predict sequences of values (time steps) by consuming their own output of the previous time step as input at subsequent time steps. In contrast to a traditional LSTM, in a ConvLSTM network, all fully connected layers are replaced with convolutional layers. We resorted to purely deterministic variants of these architectures. Models were implemented using the deep learning framework *PyTorch Lightning* [23] *which is built on top of PyTorch* [24] *and enables improved scalability. The hyperparameters were tuned using an Optuna*-based hyperparameter optimization procedure.

#### SGConvLSTM

The first deep learning architecture we tested is a ConvLSTM inspired by ref. [15]. It is termed here SGConvLSTM to reflect aspects related to the strongly guided (SG) modelling task (see above). The model is composed of *L* cells, stacked vertically. Each cell receives as input the hidden state (**h**) and memory (**c**) and an input *x* from the previous layer. Then, it outputs the updated **h***′* and **c***′*.

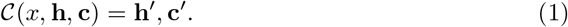

The underlying formula for a single cell is:

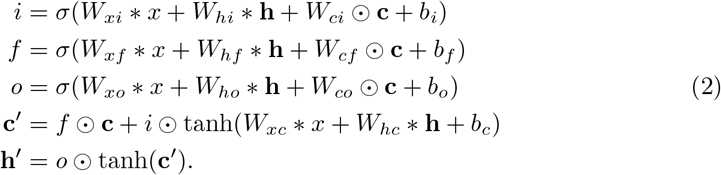

Here, *W* are the weights of the function, ∗ indicates convolution, and ⊙ the Hadamard product. Note that ∗ differs from matrix multiplication, which is used in standard LSTM [22]. Finally, the *L* cells are stacked together as in a multilayer LSTM and we take only the **h** from the deepest cell as our output. The model implemented here consists of 3 layers, where the first two layers’ cells output 20 channels and the last layer cell outputs 4 channels, corresponding to the four spectral bands (RGBI) of predicted surface reflectance. In all the convolutions, we use a kernel size of 5 × 5.

#### SGEDConvLSTM

The second model we tested was an Encoder-Decoder (ED) architecture, here referred to as SGEDConvLSTM. The Encoder-Decoder consists of two multilayer LSTM networks, as described by ref. [15]. This idea acts orthogonally to the depth of an LSTM by feeding the sequential output (sequential in depth, not in time) of the first network, the encoder, as input to a second network, the decoder, at each time step. We note that the decoder is required to have the same depth as the encoder in this setting. In such a manner, we add another dimension of parameterization to the network without having to resort solely to stacking LSTM cells on top of each other. For the SGEDConvLSTM, we mostly used the same hyperparameters as for the SGConvLSTM, except for the number of hidden channels, which was increased to 22.

#### Baseline framework

We started by using a vanilla model, i.e., a model, which is required to learn the complete image from scratch. We noticed, however, that this renders the learning process much slower. In order to leverage the peculiarity of the task (relatively small changes between subsequent images, but larger variations within images), we enhanced the model with a *baseline* (Fig. 2). The baseline was inspired by approaches such as residual connections [27] and offset regression [28]. The core idea is that our model does not need to forecast the full satellite image at the next time step, but rather only the *change* to the image, relative to the previous time step. The full next image is then computed as the sum of the previous image and the predicted change. In this manner, the model can focus on detecting how weather impacts surface reflectance changes in a given scene. We refer to embedding a predictive model such as a neural network into this general procedure as *Baseline Framework*.

**Figure 2.**
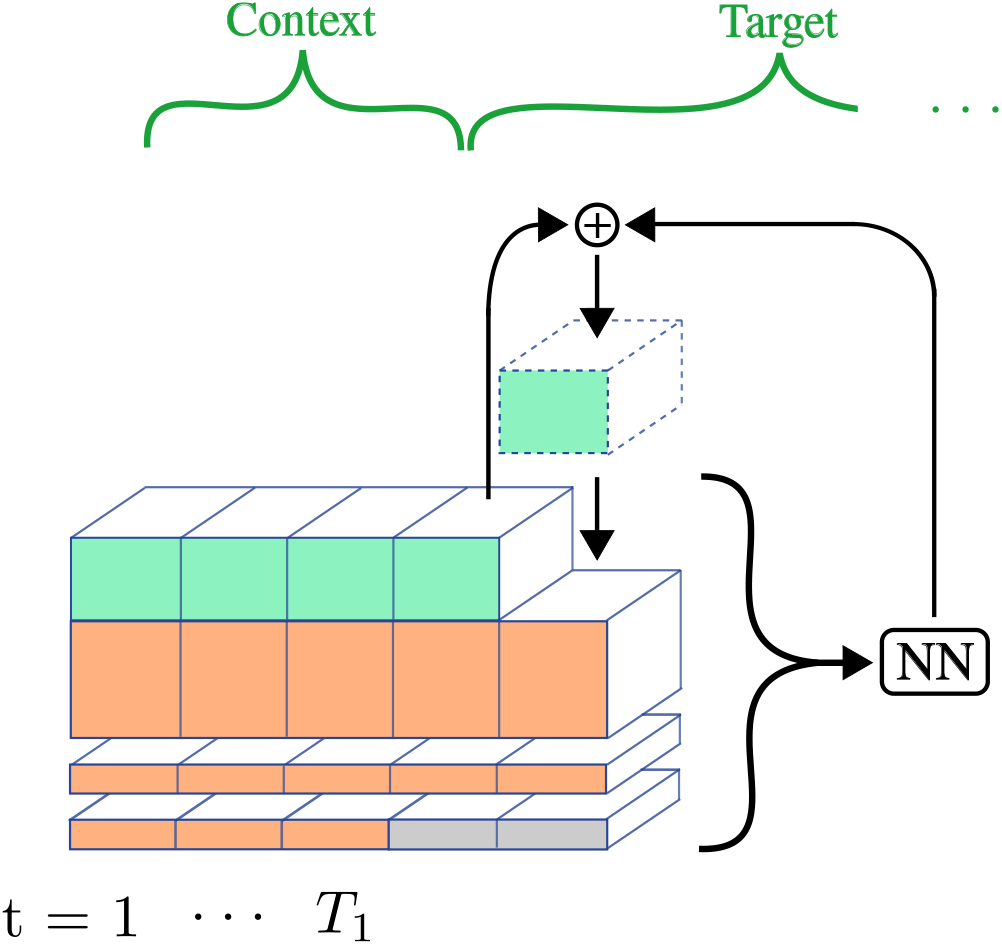
Enhancement of the recurrent model with a last frame baseline. The model predicts the change compared to the previous time step (the arrow coming from the right into the ‘+’) which are added (the ‘+’ symbol) to the previous RGBI values.

We explored different definitions of the baseline. First, we tested a baseline defined as the pixel-wise arithmetic mean across all context and previously predicted frames. This baseline is similar to the *persistence baseline* model, published by ref. [17] as a “null model” benchmark. They defined it as the pixel-wise mean across all context frames. The persistence baseline is strongly affected by outdated information from the context frames. When making predictions for future time steps, much of this data becomes irrelevant, as the images undergo significant change throughout the time steps of the target frames (multiple months). Based on this finding, we chose to use the last image of the context as the baseline for the final model. For the baseline definition based on the last image, we addressed cloud contamination in the last context frame by pixel-wise replacement with data from the last available frame in which the respective pixel was not cloud-covered. For the mean, we merely took the mean of the frames which were not cloud contaminated. Due to its improved performance (as indicated by exploratory modelling, now shown) we henceforth show results only based on the “last image baseline”.

#### Model training

23401 datacubes (97.9% of the original EarthNet2021 training dataset) were used for training, 500 datacubes for validation, and three datacubes were discarded (see above). The model was trained using the *L2 loss* determined on the predicted and observed RGBI channels (ignoring the cloud-contaminated pixels). The EarthNet score (described in Sec. 2.4) is used for validation. The learning rate is set to 0.0003 and the batch size is set to 4. We opted to use the *AdamW* optimizer, which, unlike the standard Adam optimizer [29], decouples the weight decay and has also shown to improve on generalization [30]. The SGConvLSTM and the SGEDConvLSTM were trained for 92 and 45 epochs, respectively. For completeness, a full list of the model parameters is provided in Tabs. S1 and S2 in the Supporting information.

### 2.4 Evaluation

We evaluated models following three different approaches. First, we computed the *EarthNetScore* (ENS) values, defined by ref. [17] and compared them to current entries on the EarthNet2021 leaderboard. The ENS is defined as a harmonic mean of four components, measuring complementary aspects of model performance.

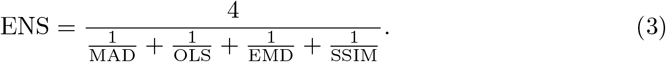

MAD is the Mean Absolute Distance. OLS is the Ordinary Least Squares (also known as L2 loss). EMD is the Earth Mover’s Distance, also known as *Wasserstein Distance*, and measures the integrated displacement of the distributions of observed and predicted values. SSIM is the Structural Similarity Index Measure and assesses the similarity of structural information in the prediction and observation, mimicking human perception of image similarity [31]. EMD and OLS are computed based on the observed and predicted Normalized Difference Vegetation Index (NDVI). MAD and SSIM are computed on all RGBI bands. The NDVI is defined based on reflectance in the red (R) and near-infrared bands (I) [32] and is computed as

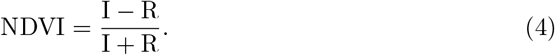

The ENS is calculated on four separate test sets, measuring different aspects of model generalizability. The independent, identically distributed (*iid*) set refers to test datacubes covering the same locations as the training set, but taken from different time intervals. The out-of-domain (*ood*) set refers to datacubes covering locations that were not part of the training set. The *extreme* test set covers locations affected by the 2018 summer drought in Central Europe (here the number of context and target frame is increased to 20 and 40, respectively). Lastly, the *seasonal* test set is similar to the *iid* set, but extending the prediction time span to approximately two years (70 context, 140 target frames). We compare our results to the EarthNet2021 scores of the initially published models *Channel-U-Net* and *Arcon* [17] (*https://www.earthnet.tech/docs/ch-leaderboard/*, last visited 3.8.2022). More recently published results by ref. [33] are used for comparison in the discussion section (Sec. 4).

For the second evaluation approach, we considered a single representative example datacube from a drought-affected scene and year (2018), taken from the extreme evaluation set. The scene is located in Saxony (Germany) and the datacube spans dates from January to November 2018, covering a reported summer drought [34, 6]. This is to visually assess model performance with a focus on the model’s ability to capture drought impacts and to gain a more intuitive understanding of different aspects of model performance than measured by aggregate metrics. In addition to the visual inspection, we examined the predicted and observed scene-average NDVI over the course of several months in summer, derived from the red and near-infrared channels of the remotely sensed surface reflectance.

Third, using the same datacube from the *extreme* evaluation set, we evaluated the predicted NDVI from a model that is forced by replaced climate data from a non-drought year (2019), and thus generates a counterfactual prediction. The rationale behind this is to assess the model’s ability to learn the climate-vegetation greenness links under anomalous conditions and predict extreme event impacts as a function of extreme climate.

## 3 Results

### 3.1 Model training efficiency

Model training on our hardware (NVIDIA GTX 1080 GPU) took ≈ 460 h. The optimization was stopped once the validation score (ENS) repeatedly failed to improve compared to the score evaluated from previous epochs. The model at the epoch with the highest attained validation score was selected. We noted a significant acceleration of convergence when employing the baseline framework, as shown in Fig. 3. The SGConvLSTM model with the last frame as the baseline achieves a validation score of 0.31 already at epoch 22. In contrast, without using the baseline, the model requires an additional 8 epochs to match this score. This illustrates that predicting a deviation on top of a specified baseline renders a simpler task than predicting the scene *ab initio*.

**Figure 3.**
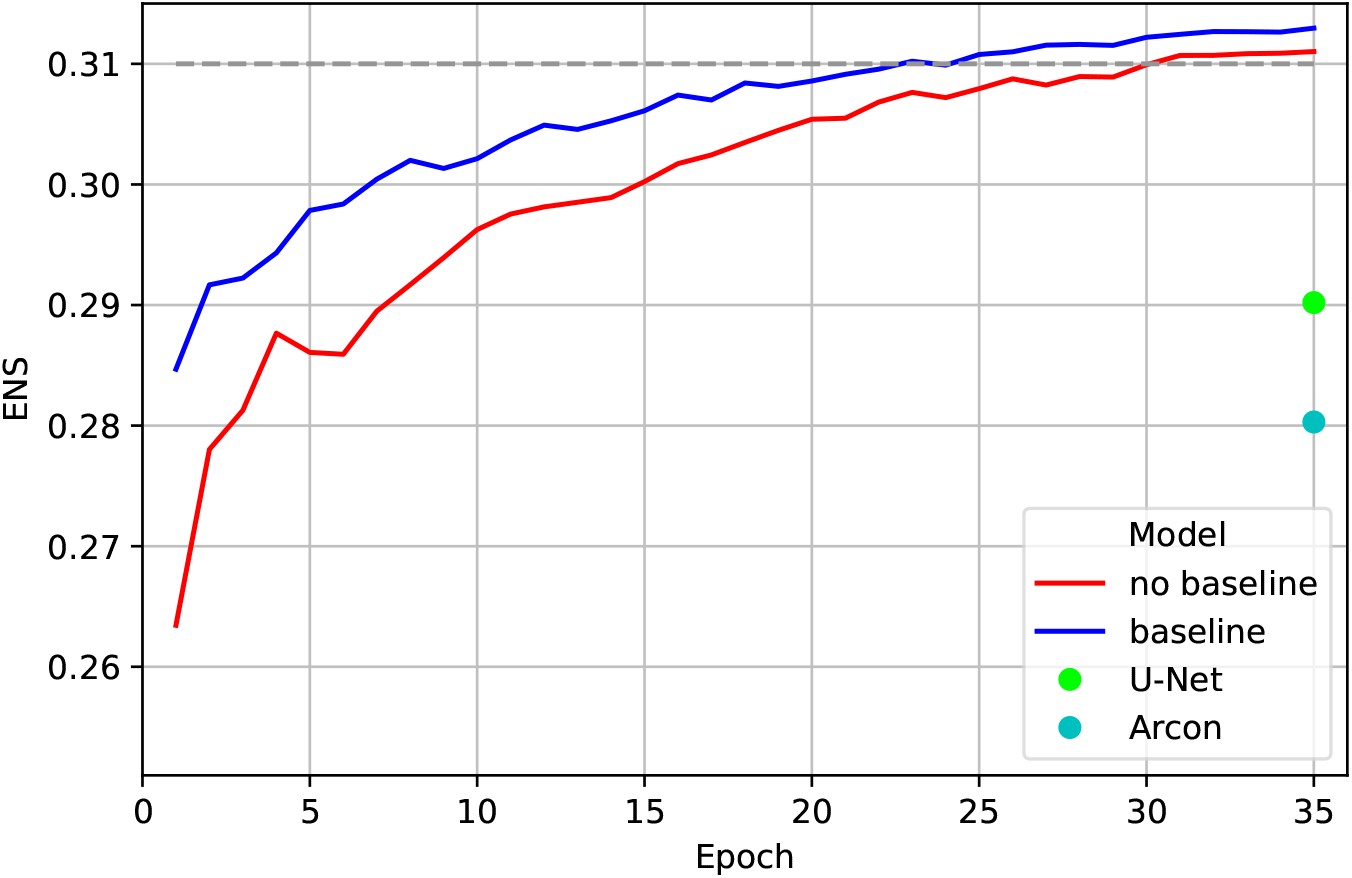
Evolution of the EarthNetScore (ENS) for increasing SGConvLSTM model training epochs, comparing alternative model setups with and without a specified baseline. For comparison, the ENS of published results by initial benchmark models (Channel-U-Net and Arcon) are plotted as dots.

### 3.2 EarthNet Score benchmarking

Both, the SGConvLSTM and the SGEDConvLSTM models show improved performance in comparison to the persistence baseline and the published results of the Channel-U-Net and Arcon models (Tab. 1 and Fig. 4). SGConvLSTM achieves an ENS of 0.3176 (an improvement of 0.0551 against the persistence baseline) on the *iid* set and outperforms the previous best model, Channel-U-Net, which achieves 0.2902 here (0.0277 better than the persistence baseline). SGEDConvLSTM achieves an ENS of 0.3164 on the *iid* set. In other words, our models’ improvements over the persistence baseline are roughly twice the improvement of Channel-U-Net.

**Table 1.** Comparison of the ENS on the four different test tracks (*iid, ood, extreme* and *seasonal*) of our models (SGConvLSTM and SGEDConvLSTM, in bold), Channel-U-Net, Arcon and Diaconu models, and the persistence baseline.

| Test set | IID | OOD | Extreme | Seasonal |
| --- | --- | --- | --- | --- |
| Persistence baseline | 0.2625 | 0.2587 | 0.1939 | 0.2676 |
| Channel-U-Net | 0.2902 | 0.2854 | 0.2364 | 0.1955 |
| Arcon | 0.2803 | 0.2655 | 0.2215 | 0.1587 |
| Diaconu | 0.3266 | 0.3204 | 0.2140 | 0.2193 |
| **SGConvLSTM** | **0.3176** | **0.3146** | **0.2740** | **0.2162** |
| **SGEDConvLSTM** | **0.3164** | **0.3121** | **0.2595** | **0.1790** |

**Figure 4.**
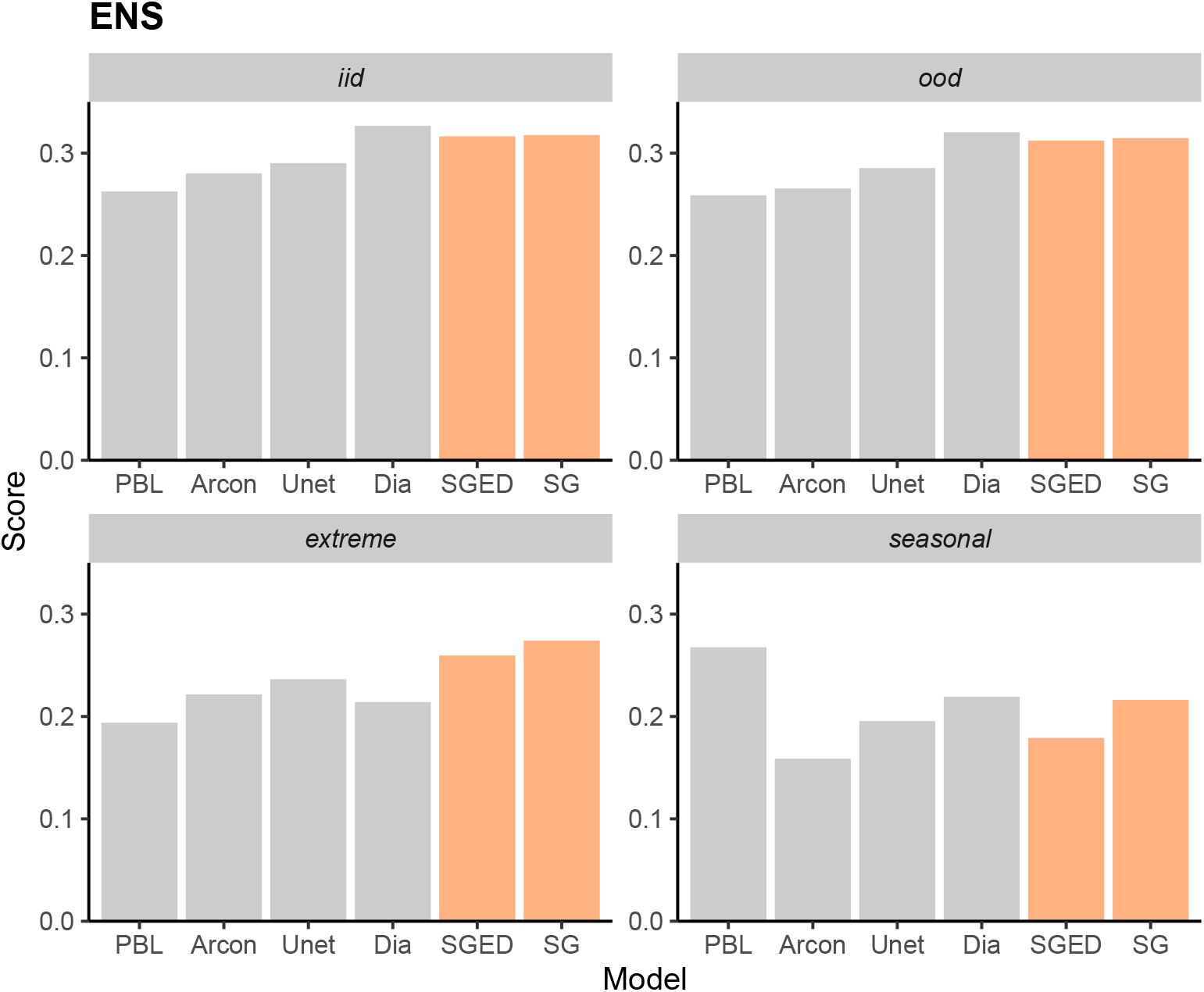
Visual comparison of the ENS on the four different test tracks (*iid, ood, extreme* and *seasonal*) of our models (’SG’ for SGConvLSTM and ‘SGED’ for SGEDConvLSTM), published Channel-U-Net (’Unet’), Arcon [17], and Diaconu et al. (’Dia’) models [33], and the published persistence baseline (’PBL’). Results from models developed here are highlighted in orange color.

The negligible differences between the *iid* and *ood* scores of the SGConvLSTM and SGEDConvLSTM models underline the models’ ability to generalize well to data outside the training set. Here, our models’ score degrades only by 0.003 and 0.004, respectively. In contrast, the score of Arcon declines by 0.015, thus showing a poorer out-of-training-distribution generalization capability.

Overall, the performance of SGConvLSTM is slightly better than the performance of SGEDConvLSTM, both on the *iid* and *ood* test sets, where it attained 0.3176 and 0.3146, respectively, compared to an ENS of 0.3164 and 0.3121 achieved by SGEDConvLSTM for the *iid* and *ood* test sets.

The strength of models developed here is most evident for the evaluation using the *extreme* test set. The SGConvLSTM (SGEDConvLSTM) reached an ENS of 0.2740 (0.2595), an increase of 0.080 (0.066) compared to the persistence baseline. In contrast, Channel-U-Net only improves by 0.043 over the persistence baseline. This suggests that our models provide more informative predictions of the development of vegetation greenness under future extreme climatic conditions than current benchmarks.

The evaluation on the *seasonal* test set revealed generally weaker model performance compared to performances on the other test sets. Neither of our models outperformed the persistence baseline. This suggests that models presented here, as well as the other published benchmarks, provide less reliable predictions at the seasonal-to-annual timescale (here 140 frames, corresponding to roughly two years).

Evaluating component metrics of the ENS score (Tab. S3 and Figs. S4-S7) reveals additional information about the robustness of different models. Large differences in model performance are evident in particular for the structural similarity metric (SSIM) and for all metrics when comparing model performance on the *extreme* and the *seasonal* test sets with performances on the *iid* and *ood* test sets. For example, for the *extreme* test set, the models presented here (SGConvLSTM and SGEDConvLSTM) show substantial improvements over the persistence baseline and over other Channel-U-Net and Arcon models. This is most evident when considering the SSIM (Tab. S3 and Fig. S7). Lacking robustness in long-term predictions (*seasonal* test set) of all models appears to be linked in particular to the pronounced deterioration in the SSIM. In contrast, structural similarity is maintained better in the shorter-term predictions assessed by the *iid* and *ood* test tracks.

### 3.3 Example scene analysis

The evaluation of the example scene (Figs. 5 and Fig. S1) reveals that models predict the distinct evolution of vegetation greenness across different portions of the image, representing different landscape elements, land cover, vegetation types, and individual fields. However, although structural patterns are reliably modelled, distinct greenness changes in different fields at different points in time appear to be outstanding challenges for the models assessed here.

**Figure 5.**
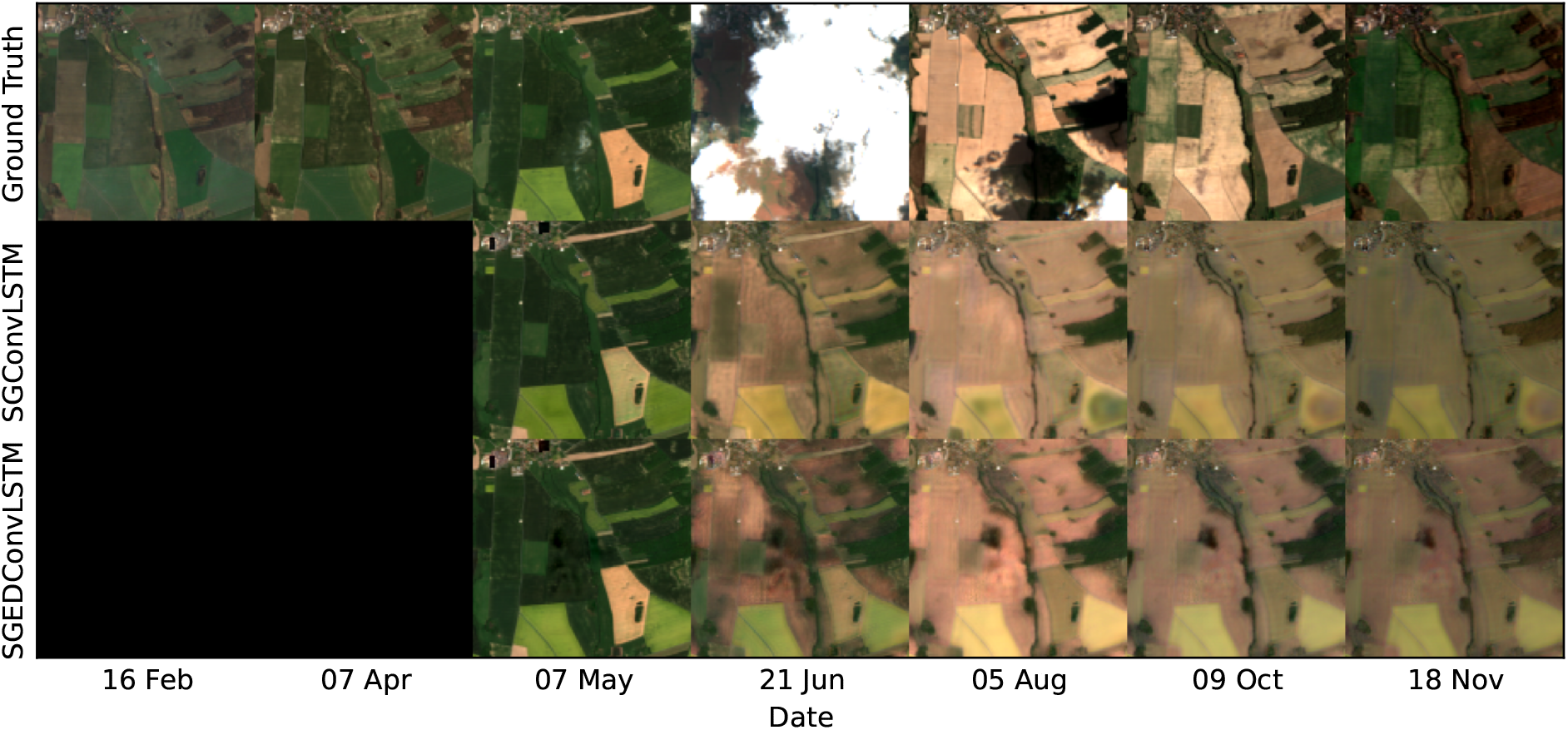
Observed (top row) and modelled (two bottom rows for the SGEDConvLSTM and the SGConvLSTM models) real-color images of the surface reflectance in RGB channels for an example scene from the *extreme* test set, located in Saxony, Germany, and covering dates from February to November 2018. Data is shown for roughly evenly distanced time steps, avoiding images with clouds where possible. The first two columns are in the context section and are not forecasted by the models. Model predictions are for 7 May and subsequent time steps.

To further assess the reliability of our models, we provide as Supporting information an equivalent image for a datacube in the *iid* dataset Fig. S2 and the *ood* dataset Fig. S3. No clear differences in prediction accuracy are evident from visual comparison of the scenes taken from the *iid* and the *ood* test tracks, reflecting also the similar ENS score achieved on the *iid* and the *ood* test tracks (Tabs. 1 and S3). This corroborates the out-of-distribution generalization properties of our models, identified above (Sec. 3.2).

### 3.4 Counterfactual analysis

Results for the SGConvLSTM model show substantially different simulated responses of surface reflectance when using climate forcing data from a drought-affected year (2018) versus data from a year without drought in the respective location (2019) (Figs. 6 and 7). The visual comparison of RGB images (Fig. 6) indicates unrealistic, excessively green vegetation when the model is forced by counterfactual climate, taken for corresponding days and months from year 2019. However, this visualisation also indicates an overly sensitive simulated response (excessive browning) when the model is forced by actual weather. When aggregating the mean NDVI across the same scene and evaluating the temporal course of observed and modelled NDVI (Fig. 7), we find that the onset of browning (i.e., decline of the NDVI after its seasonal maximum) is simulated roughly half a month too early for both models (SGConvLSTM and SGEDConvLSTM), but the NDVI attains similar levels in predictions and observations around one month after the onset of browning. In contrast, when models are forced by (counterfactual) 2019 climate, the NDVI remains too high compared to observations throughout the period assessed.

**Figure 6.**
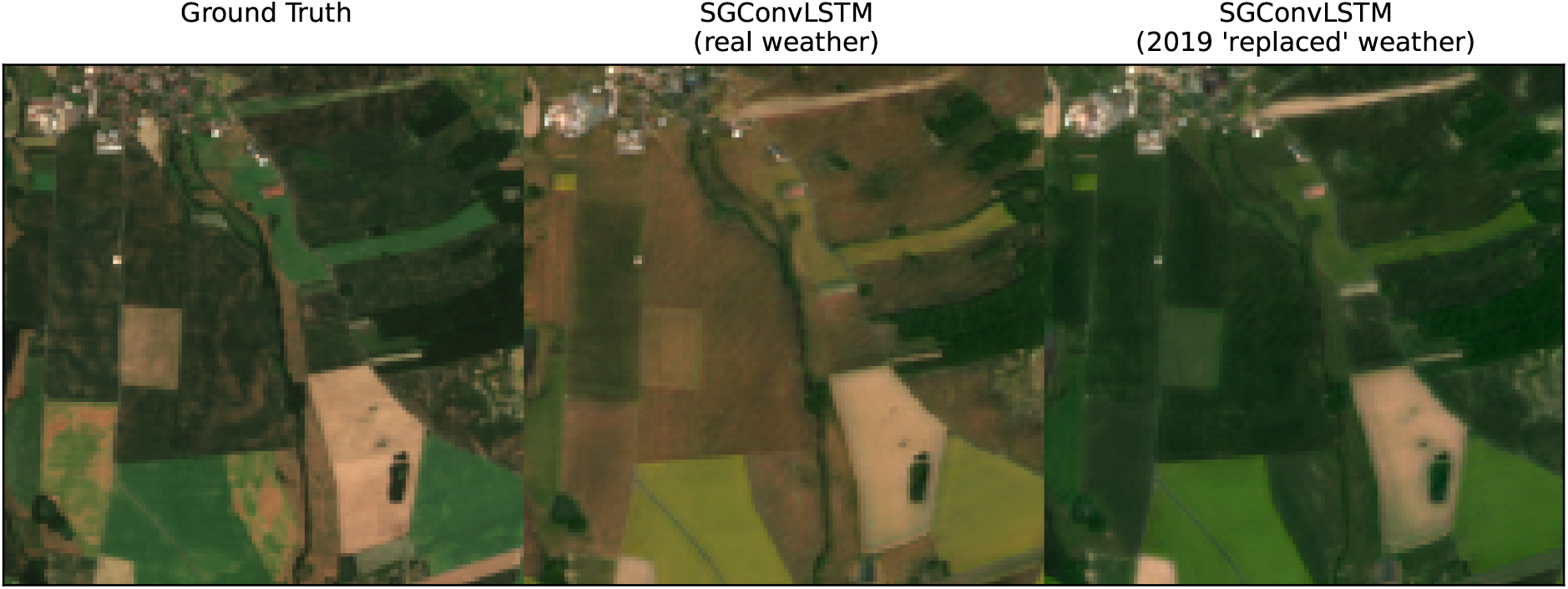
(Left) Ground truth, (Middle) forecast satellite image using 2018 weather and (Right) forecasted satellite image using 2019 weather in Saxony (Germany). All images correspond to the 5th of June, the first day when significant browning is observed in 2018, but not in 2019. This day corresponds to the vertical dotted line in Fig. 7.

**Figure 7.**
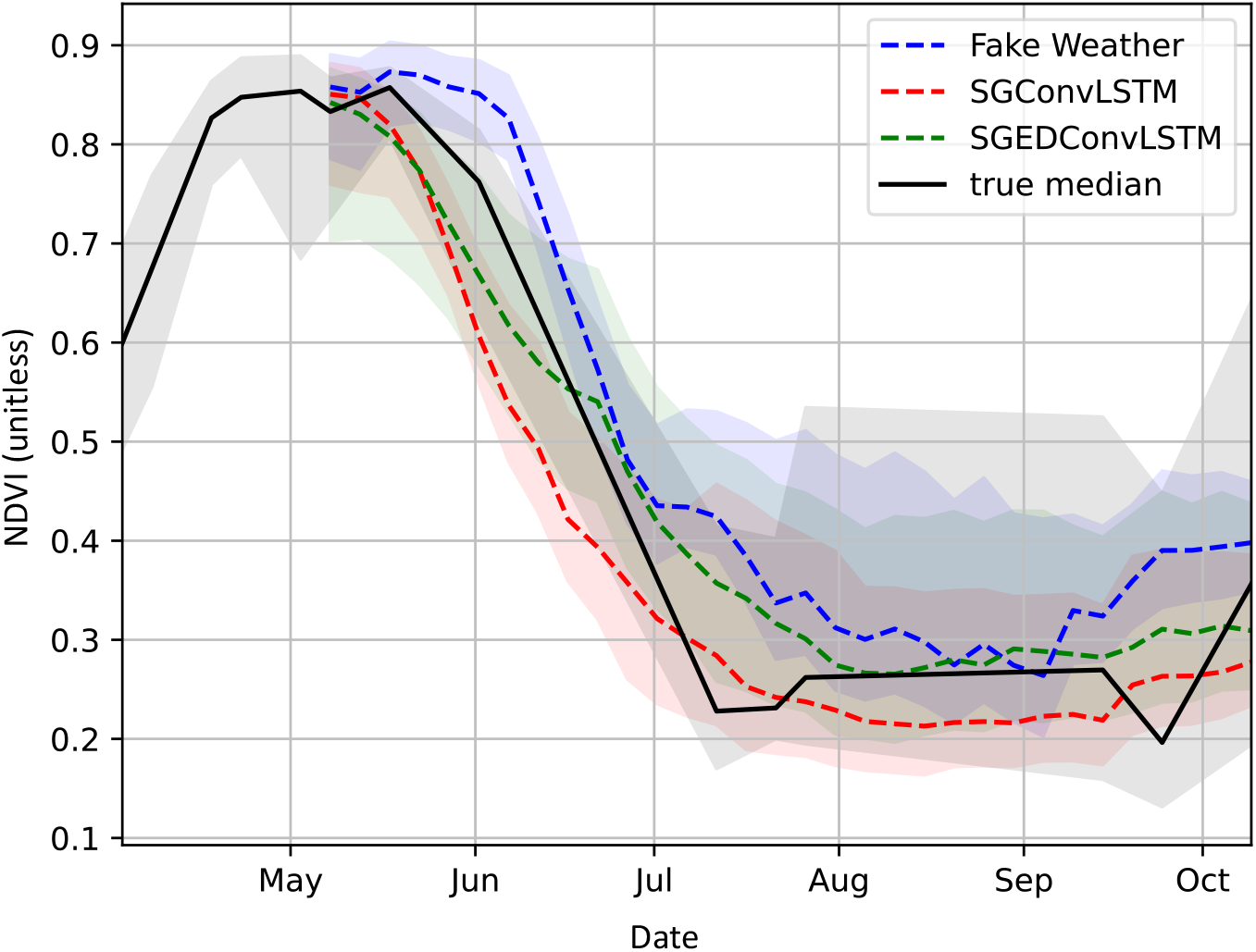
Time series of the NDVI within a specific datacube in Saxony (Germany) in 2018. The black solid line indicates the median NDVI, the shaded area indicates the region between the 0.25 quantile and the 0.75 quantile. Note that the ground truth is subject to some missing NDVI data points: an artifact that originates from the presence of clouds. This leads to the impossibility of computing the true NDVI for some time steps. We address this issue by means of linear interpolation using available NDVI from neighboring time steps. The 2019 weather refers to the experiment where we utilize 2019 weather features from the same location.

### 4 Discussion

Drought stress limits vegetation activity across a large portion of the Earth’s land surface [35, 36, 37], and, under extreme conditions, impacts land surface greenness [2], ecosystem productivity [38], and plant health [39] and also in relatively moist regions. Although retrospective analyses of remote sensing data allows an identification of the timings and locations of discernible impacts of drought stress on surface reflectance and vegetation greenness, only few studies have used these observations in combination with data on environmental covariates to establish functional relationships and develop predictive data-driven models (but see ref. [33]). Here, we developed deep learning models that combine convolutional layers and LSTM, thus making use of the spatio-temporal dependencies in the data.

### 4.1 Comparison to published models

Following the standardized EarthNet2021 evaluation protocol [17], we show that our models clearly outcompete a “null model” of a pixel-wise constant mean surface reflectance and greenness (Persistence Baseline), and perform better that initially published models (Channel-U-Net and Arcon [17]). Using additional analyses of an example scene from a location and year that is known to have been affected by a summer drought (2018 in Saxony, Germany), we demonstrate that the models presented here make use of climate information to predict vegetation greenness and that models predict anomalous land surface browning under anomalously dry conditions. The demonstrated model skill (relative to the “null model”), assessed on out-of-sample scenes (*ood* test track) further demonstrates the capability of our models to generalise across space and to predict the evolution of surface reflectance at sites for which data has not been used during model training. In other words, the models have potential to scale vegetation greenness forecasts across space.

Our models show similar performance compared to recently published results by ref. [33] who used a similar model (also a ConvLSTM), but with a different specification of the target (not using the baseline framework as applied here). Following the *extremes* evaluation track, models presented here exhibit improved performance over the model presented by ref. [33]. Model performance on the *extremes* evaluation track is particularly relevant in the context of early warning of forest damage or agricultural yield loss as a consequence of drought conditions. However, the model presented by ref. [33] appears to suffer less from longer-term “drift” of the predicted distributions, compared to the models presented here - as shown by comparing the EMD metric on the seasonal test track.

Following the approach followed here (*baseline framework*), we used an initial baseline for prediction, and target only the incremental deviations (*delta*) of the subsequent time steps from the baseline. We chose this instead of directly predicting the full RGBI features, as it rendered improved model training efficiency and final model performance. This approach is neither specific to a particular underlying neural network architecture (ConvLSTM or Encoder-Decoder ConvLSTM), nor to a specific choice of the baseline, but comes with some limitations. We note that despite the significant short-term performance gains of our models when evaluating on the *iid, ood* and the *extreme* datasets, long-term predictions were poor compared to the simple persistence baseline. This is likely due to the additive nature of error accumulation, and to the fact that model training was performed on a much shorter target window compared to the target sequence in the *seasonal* test track, and thus did not account for the full seasonal cycle reflected in the latter.

### 4.2 Prediction challenge specification

Data-driven drought impact prediction is in its infancy. The EarthNet2021 Challenge fosters development of research in this direction and the present study is an attempt at satellite image forecasting that yielded several insights to guide future work. Reliable early warning for stakeholders requires accurate predictions of greenness changes for different portions of the image, representing different land cover and land use classes. E.g., forest managers require information of expected impacts on forest greenness - a proxy for tree vitality. The task as defined here includes greenness predictions in cropland areas. There, crop phenophases and dates of sowing, harvesting, and ploughing strongly affect surface reflectance - the prediction target - and are thus implicitly part of the prediction task. However, surface changes in cropland areas are subject to deliberate decisions by the farmer and are affected by differences between crop types and cultivars. A possibility is that future iterations of the EarthNet2021 Challenge specify the prediction task to be more directly tailored to potential applications and stakeholder interests and reduce the scope of evaluated predictions to corresponding land cover classes.

Given the specification of the EarthNet2021 Challenge and its data, the prediction target is likely dominated by structural aspects (large variations in surface reflectance within a scene), and seasonal variations (20-140 target frames) in surface reflectance and vegetation greenness. Slighter nuances in greenness within portions of the image and distinct sensitivity within land cover types and within individual agricultural fields constitute a smaller fraction of the overall variation of the data and are thus likely treated by models as “second-order effects”. However, these nuances bear very relevant information for process understanding, linked, e.g., to topographic effects that modify climate impacts across the landscape [10]. Future work may develop models that reduce the scope of the prediction task to learn these nuances and thus learn about heterogeneity of climate impacts, depending on the topographic position, and (if sufficiently high-quality and -resolution data is available) physical soil and bedrock properties. Establishing these relationships will be important for addressing open research challenges in ecohydrology [10] and may have to rely on methods of *explainable machine learning*. By following the *baseline framework*, we implemented such a “scope reduction” by targeting only the deviation of the surface reflectance over time from an initial baseline. This thus emphasizes the drought-related browning of vegetation in summer, while the baseline with its large spatial variations within a scene is provided as the initial state.

### 4.3 Methodological advances

Initial approaches to the EarthNet2021 challenge use existing video prediction solutions [17]. The Channel-U-Net model uses an architecture roughly based on *U-Net* [40], where all input data is stacked along the channel dimension and does not model the temporal dependencies explicitly. The second model, Arcon, is an adaptation of the Stochastic Adversarial Video Prediction (SAVP) [41] model which does model temporal dependencies, but it was designed primarily for a highly stochastic setting (including moving objects) in the general video prediction context. Both types of models appear to be less well suited as deep learning solutions for satellite image predictions.

Various extensions to the ConvLSTM for sequential image prediction have been proposed, leaving room for future methodological improvements. These include ensembling multiple ConvLSTMs to better tackle the (in our case ecological and land use) diversity of the data [42]. In order to “encourage” the model to focus on noticeable spatial features, attention mechanisms such as soft attention [43] have also been integrated into ConvLSTMs successfully [44]. However, we also noticed that the models trained here exhibited no tendency to overfit and the training data volume may be further increased by enlarging the sample of datacubes. Therefore, we expect that further gains can be achieved without resorting to different model architectures, but by developing architectures with higher parameterization, e.g., by adding more layers, increasing the dimensionality of hidden channels, or increasing the kernel size.

Recurrent architectures are not the only means for capturing time dependencies effectively. In recent years, Transformer-based architectures [45] have led to remarkable successes in numerous applications - besides natural language processing [46, 47, 48]. Since these are not conceived in a sequential manner, they exhibit multiple advantages over recurrent architectures, including a more direct gradient flow, a higher level of parallelizability [49] and allowing for effective self-supervised pre-training schemes [50].

In our efforts to use a Transformer version for video prediction, called *ConvTransformer* [16], we encountered significant memory limitations, even after decreasing the hidden channel dimension and resorting only to single attention heads. In the proposed architecture, the so-called values need to be replicated many times and be kept in memory together with *keys* and *queries* all at once for efficient computation. This procedure is rendered infeasible, e.g., in the seasonal setting where we aim to predict 140 frames, using the hardware at our disposal (GPU with 12 GB memory).

## 5 Outlook and conclusion

While models are trained and evaluated here on data from the past - using observational surface reflectance and climate from reanalysis - future applications may include generating actual forecasts where drought impact models are forced by numerical weather predictions. While our evaluation of the modelled surface reflectance suggests relatively reliable predictions for twenty future frames (∼100 days), medium (15 days) to and long-range (months) weather predictions have limited reliability [51] and will therefore likely constitute a dominating source of error in an actual forecasting context.

The seasonal development of vegetation greenness is mechanistically linked to ecosystem-level photosynthesis and vegetation primary productivity [52, 53]. Combined with additional data sources and model “layers”, greenness forecasts may thus provide the basis for modelling additional targets, including agricultural yields or wood production.

In this work, we demonstrate the benefit of Convolutional LSTMs for satellite image prediction and drought response forecasting using incremental inference from a prior baseline to predict future drought responses. Our methodology shows potential of using general video prediction methods in capturing both temporal dependencies and spatial structure across the landscape in response to climate drivers and to scale predictions in space.

### 5.1 Code availability

All code is available online: https://zenodo.org/record/6985292

## 5.2 Acknowledgements

We thank Ce Zhang for initial discussions that helped us with this work. B.D.S was funded by the Swiss National Science Foundation grant PCEFP2 181115. K.H. was supported by the generosity of Eric and Wendy Schmidt by recommendation of the Schmidt Futures program.

## S1 Supporting information

**Table S1.** Hyperparameters for ConvLSTM.

| Parameter | Meaning | Value |
| --- | --- | --- |
| n_layers | Depth of the network | 3 |
| h_channel | Number of hidden channels (width of the network) | 20 |
| kernel_size | Size of the kernel | 5x5 |
| epochs | Number of training epochs | 92 |
| train_loss | Loss used for training | L2 |
| val_loss | Loss used for validation (and testing) | ENS |
| learn_rate | Learning rate | 0.0003 |
| batch_size | Number of training datacubes in a batch | 4 |
| optimizer | Optimizer used for training | AdamW |

**Table S2.**
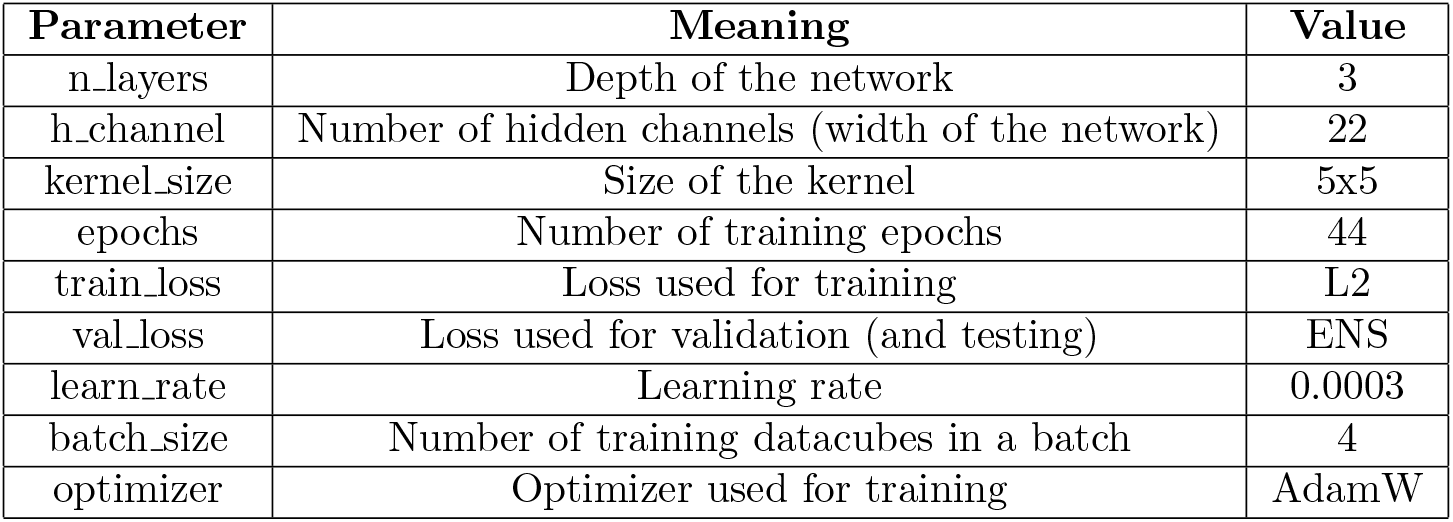
Hyperparameters for EncoderDecoderConvLSTM

| Parameter | Meaning | Value |
| --- | --- | --- |
| n_layers | Depth of the network | 3 |
| h_channel | Number of hidden channels (width of the network) | 22 |
| kernel_size | Size of the kernel | 5x5 |
| epochs | Number of training epochs | 44 |
| train_loss | Loss used for training | L2 |
| val_loss | Loss used for validation (and testing) | ENS |
| learn_rate | Learning rate | 0.0003 |
| batch_size | Number of training datacubes in a batch | 4 |
| optimizer | Optimizer used for training | AdamW |

**Table S3.** Extended ENS comparison, including the ENS components for our models and the previous state-of-the-art (Channel-U-Net and Arcon).

|  | IID |  |  |  |  |
| --- | --- | --- | --- | --- | --- |
|  | ENS | MAD | OLS | EMD | SSIM |
| Persistence baseline | 0.2625 | 0.2315 | 0.3239 | 0.2099 | 0.3265 |
| Channel-U-Net | 0.2902 | 0.2482 | 0.3381 | 0.2336 | 0.3973 |
| Arcon | 0.2803 | 0.2414 | 0.3216 | 0.2258 | 0.3863 |
| Diaconu | 0.3266 | 0.2638 | 0.3513 | 0.2623 | 0.5565 |
| **SGConvLSTM (this paper)** | **0.3176** | **0.2589** | **0.3456** | **0.2533** | **0.5292** |
| **SGEDConvLSTM (this paper)** | **0.3164** | **0.2580** | **0.3440** | **0.2532** | **0.5237** |
|  | OOD |  |  |  |  |
|  | ENS | MAD | OLS | EMD | SSIM |
| Persistence baseline | 0.2587 | 0.2248 | 0.3236 | 0.2123 | 0.3112 |
| Channel-U-Net | 0.2854 | 0.2402 | 0.3390 | 0.2371 | 0.3721 |
| Arcon | 0.2655 | 0.2314 | 0.3088 | 0.2177 | 0.3432 |
| Diaconu | 0.3204 | 0.2541 | 0.3522 | 0.2660 | 0.5125 |
| **SGConvLSTM (this paper)** | **0.3146** | **0.2512** | **0.3481** | **0.2597** | **0.4977** |
| **SGEDConvLSTM (this paper)** | **0.3121** | **0.2497** | **0.3450** | **0.2587** | **0.4887** |
|  | Extreme |  |  |  |  |
|  | ENS | MAD | OLS | EMD | SSIM |
| Persistence baseline | 0.1939 | 0.2158 | 0.2806 | 0.1614 | 0.1605 |
| Channel-U-Net | 0.2364 | 0.2286 | 0.2973 | 0.2065 | 0.2306 |
| Arcon | 0.2215 | 0.2243 | 0.2753 | 0.1975 | 0.2084 |
| Diaconu | 0.2140 | 0.2137 | 0.2906 | 0.1879 | 0.1904 |
| **SGConvLSTM (this paper)** | **0.2740** | **0.2366** | **0.3199** | **0.2279** | **0.3497** |
| **SGEDConvLSTM (this paper)** | **0.2595** | **0.2304** | **0.3164** | **0.2186** | **0.2993** |
|  | Seasonal |  |  |  |  |
|  | ENS | MAD | OLS | EMD | SSIM |
| Persistence baseline | 0.2676 | 0.2329 | 0.3848 | 0.2034 | 0.3184 |
| Channel-U-Net | 0.1955 | 0.2169 | 0.3811 | 0.1903 | 0.1255 |
| Arcon | 0.1587 | 0.2014 | 0.3788 | 0.1787 | 0.0834 |
| Diaconu | 0.2193 | 0.2146 | 0.3778 | 0.2003 | 0.1685 |
| **SGConvLSTM (this paper)** | **0.2162** | **0.2207** | **0.3756** | **0.1723** | **0.1817** |
| **SGEDConvLSTM (this paper)** | **0.1790** | **0.2056** | **0.3585** | **0.1543** | **0.1218** |

**Figure S1.**
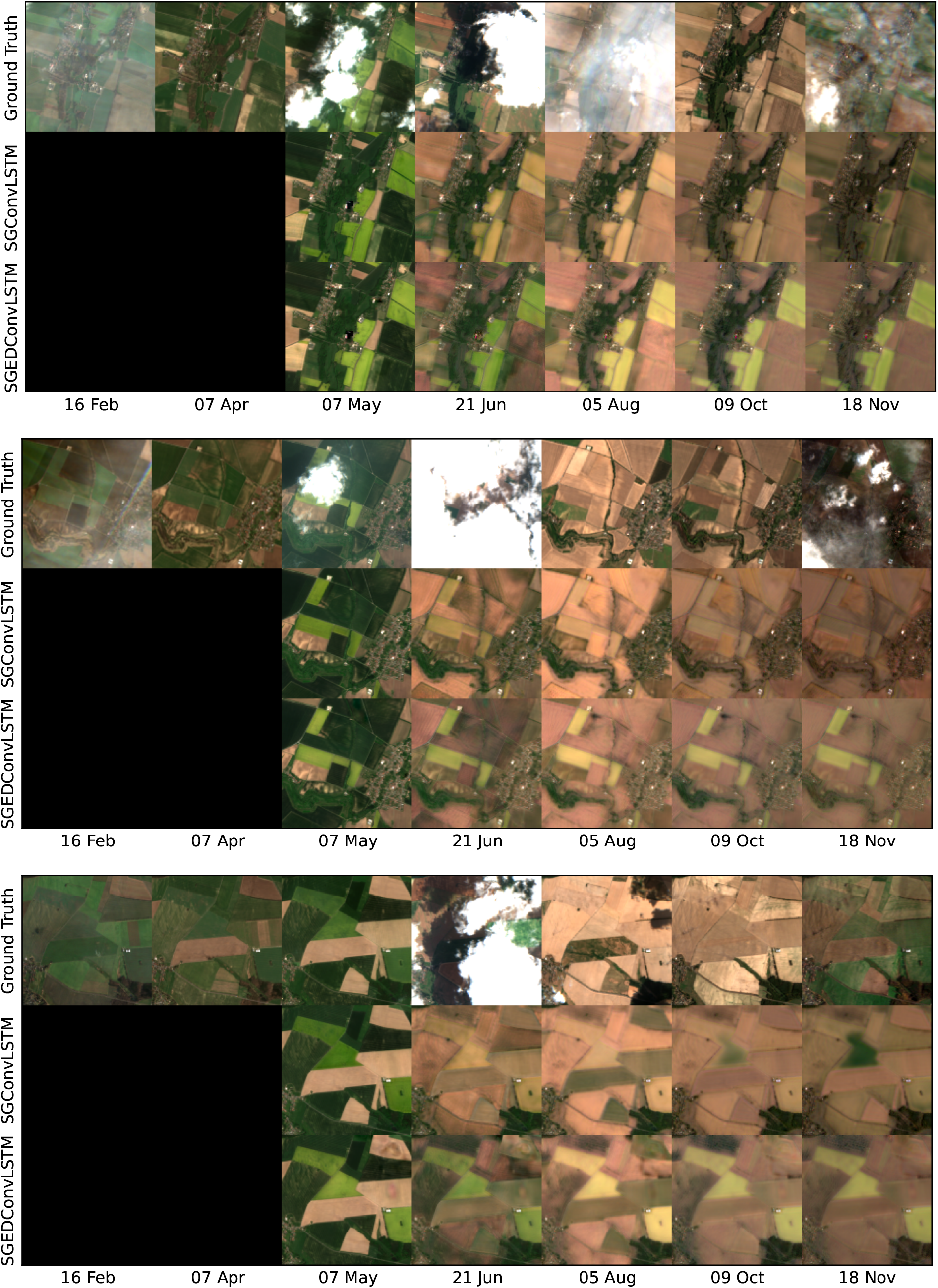
Three additional extreme datacube predictions (located in Germany in 2018).

**Figure S2.**
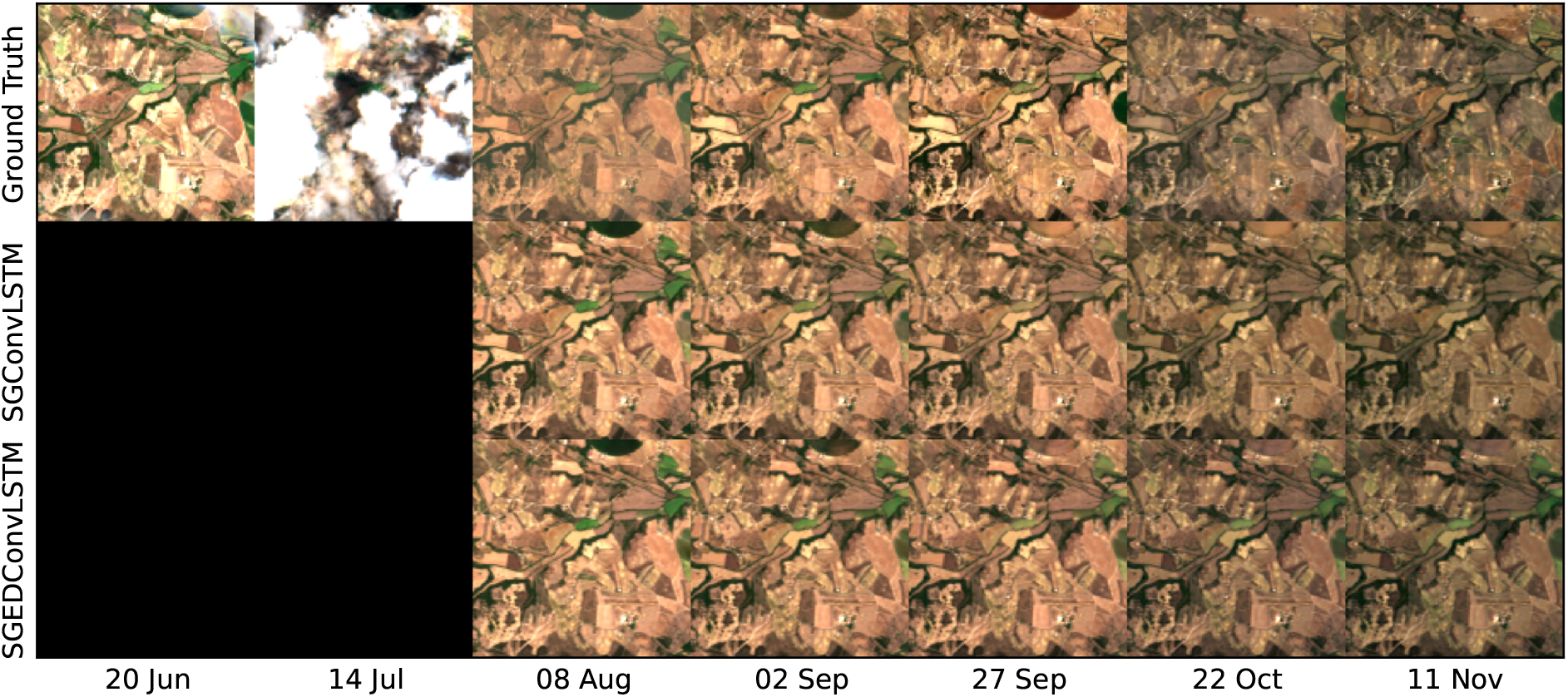
Observed (top row) and modelled (two bottom rows for the SGEDConvLSTM and the SGConvLSTM models) surface reflectance in RGB channels for an example scene from the *iid* test set, located in Portugal, and covering dates from June - November 2017. Data is shown for roughly evenly distanced time steps, avoiding images with clouds where possible. The first two columns are in the context section and are not forecasted by the models.

**Figure S3.**
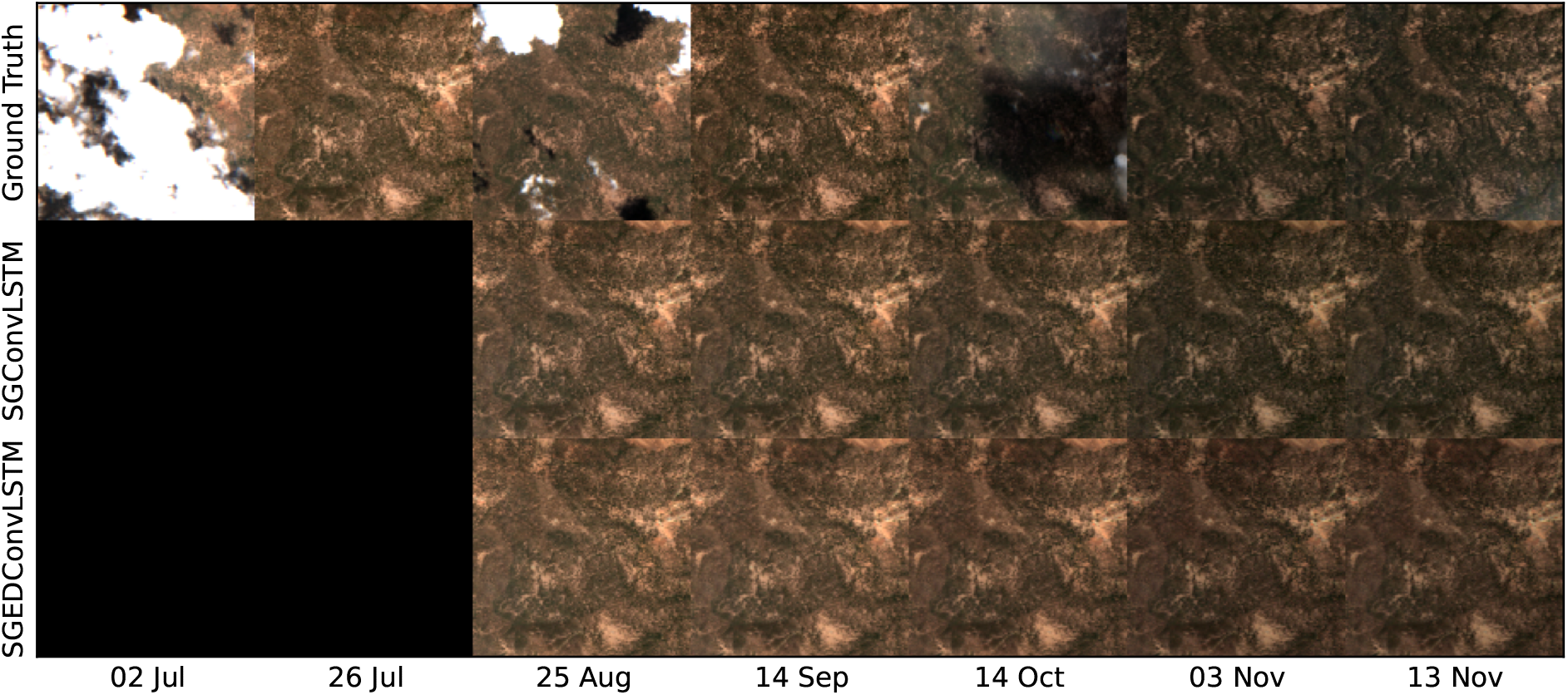
Observed (top row) and modelled (two bottom rows for the SGEDConvLSTM and the SGConvLSTM models) surface reflectance in RGB channels for an example scene from the *ood* test set, located in Andalusia, Spain, and covering dates from July - November 2017. Data is shown for roughly evenly distanced time steps, avoiding images with clouds where possible. The first two columns are in the context section and are not forecasted by the models.

**Figure S4.**
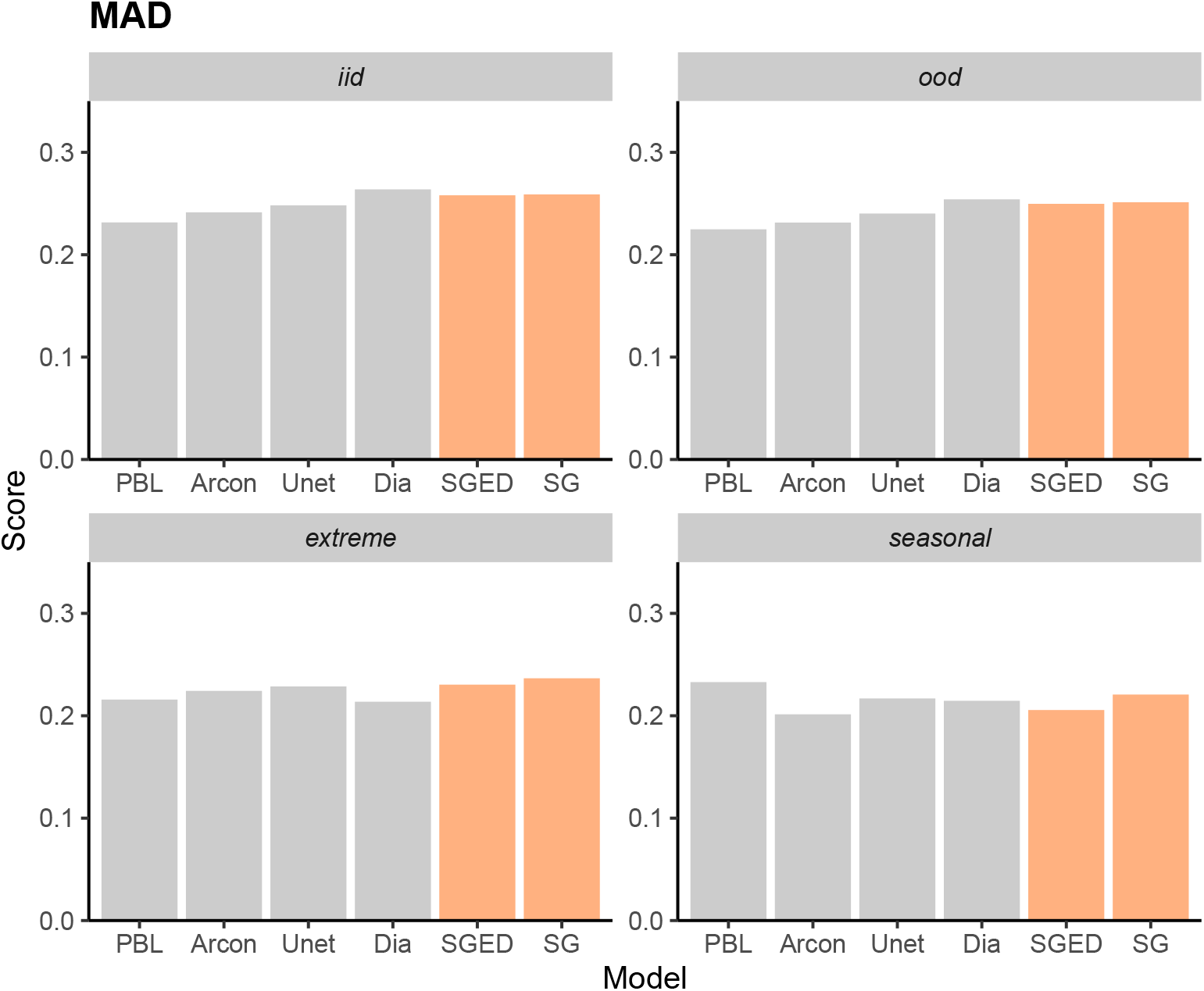
Visual comparison of the MAD on the four different test tracks (*iid, ood, extreme* and *seasonal*) of our models (’SG’ for SGConvLSTM and ‘SGED’ for SGEDConvLSTM), published Channel-U-Net (’Unet’), Arcon, and Diaconu et al. [33] (’Dia’) models, and the persistence baseline (’PBL’).

**Figure S5.**
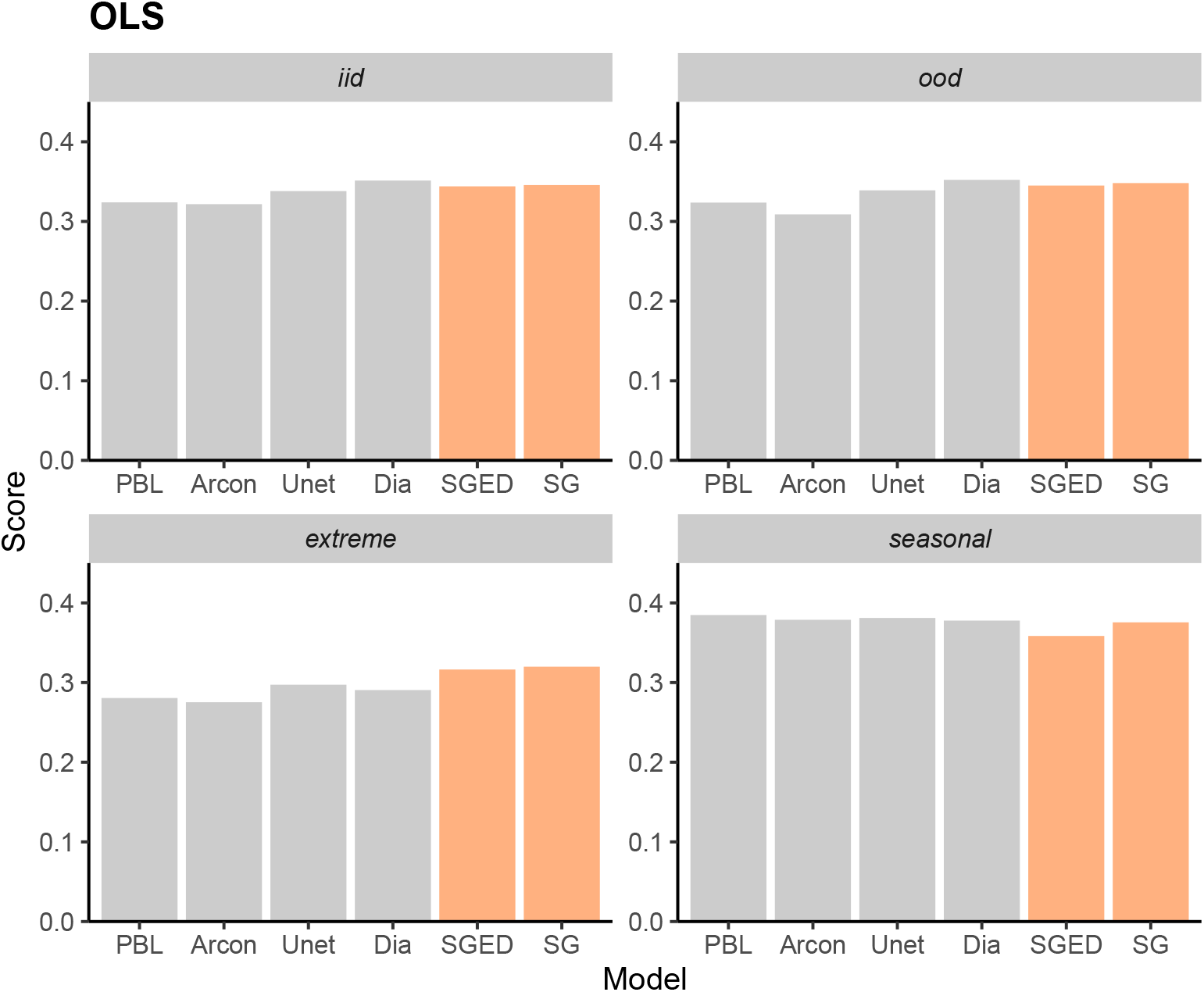
Visual comparison of the OLS on the four different test tracks (*iid, ood, extreme* and *seasonal*) of our models (’SG’ for SGConvLSTM and ‘SGED’ for SGEDConvLSTM), published Channel-U-Net (’Unet’), Arcon, and Diaconu et al. [33] (’Dia’) models, and the persistence baseline (’PBL’).

**Figure S6.**
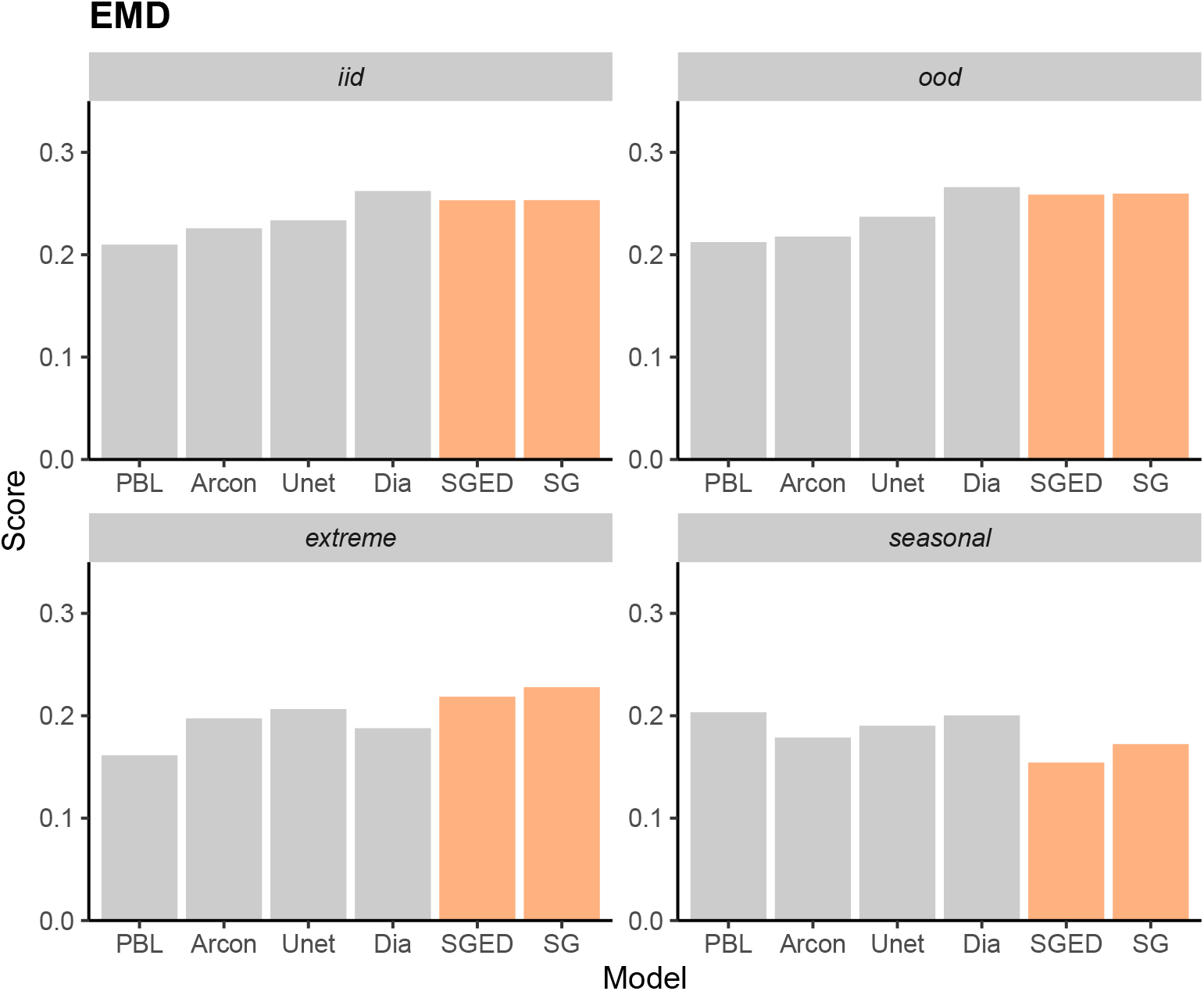
Visual comparison of the EMD on the four different test tracks (*iid, ood, extreme* and *seasonal*) of our models (’SG’ for SGConvLSTM and ‘SGED’ for SGEDConvLSTM), published Channel-U-Net (’Unet’), Arcon, and Diaconu et al. [33] (’Dia’) models, and the persistence baseline (’PBL’).

**Figure S7.**
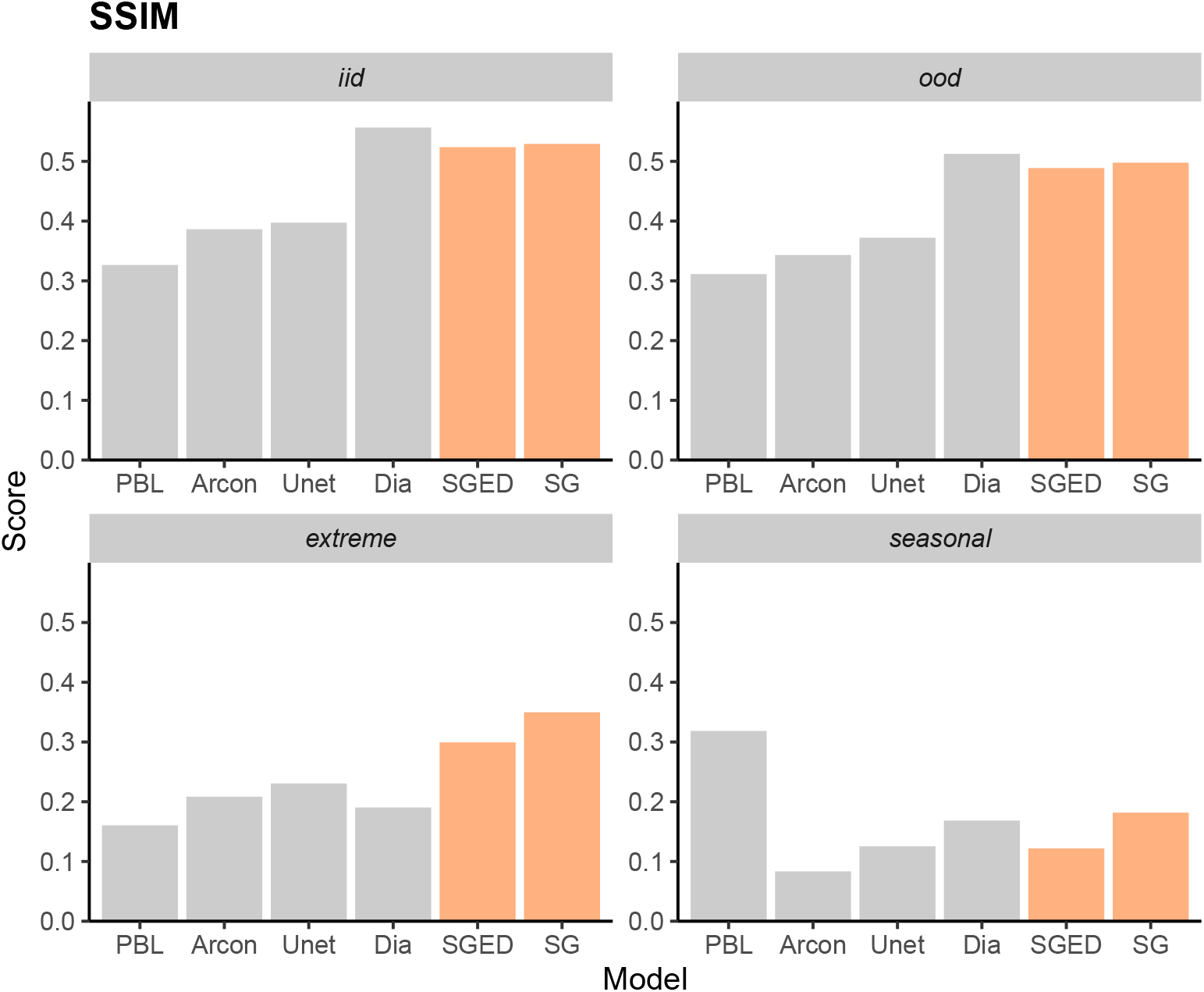
Visual comparison of the SSIM on the four different test tracks (*iid, ood, extreme* and *seasonal*) of our models (’SG’ for SGConvLSTM and ‘SGED’ for SGEDConvLSTM), published Channel-U-Net (’Unet’), Arcon, and Diaconu et al. [33] (’Dia’) models, and the persistence baseline (’PBL’).

